# Electrical Surface Polarization as a Functionalization Strategy to Improve Bone Regeneration of Apatite-Based Graft Materials

**DOI:** 10.64898/2026.02.17.705299

**Authors:** Katja Hrovat, Leire Bergara Muguruza, Rumi Hiratai, Ari Alho, Maarit Laine, Keijo Mäkelä, Kimihiro Yamashita, Miho Nakamura

**Affiliations:** Medicity Research Laboratory, Faculty of Medicine, University of Turku, Turku, Finland; Laboratory for Biomaterials and Bioengineering, Institute of Science Tokyo, Tokyo, Japan; Department of Anesthesia and Intensive Care, Turku University Hospital and University of Turku, Finland; Department of Orthopaedics and Traumatology, Turku University Hospital and University of Tuku, Turku, Finland; Graduate School of Medical and Dental Science, Institute of Science Tokyo, Tokyo, Japan; Advanced Central Research Organization, Teikyo University, Tokyo, Japan; La Salle Campus Barcelona Ramon Llull University, Barcelona, Spain

## Abstract

Apatite-based bone graft materials are widely used for bone regeneration; however, their limited bioactivity and slow remodeling often hinder complete replacement by newly formed bone. Electrical surface polarization has emerged as a promising non-chemical strategy to modify biomaterial surface properties without altering bulk characteristics. In this study, we investigated the effects of electrical surface polarization on apatite-based biomaterials using synthesized carbonate apatite (CA) for mechanistic in vitro evaluation and a clinically relevant xenograft material for in vivo validation.

Material characterization confirmed the formation of B-type carbonate apatite with bone-like mineral composition. Thermally stimulated depolarization current measurements verified successful induction of surface charges, with polarization intensity dependent on treatment conditions. In vitro studies using human peripheral blood–derived osteoclast precursors demonstrated that electrically polarized CA surfaces significantly enhanced osteoclast differentiation and resorptive activity compared to non-polarized controls, with the strongest effects observed on positively polarized surfaces. Three-dimensional analysis revealed increased resorption pit depth and volume, indicating enhanced osteoclast functionality.

In vivo implantation of polarized xenograft materials into rat femoral defects resulted in significantly increased new bone formation and improved implant–bone integration compared to non-polarized materials. Higher polarization conditions promoted more mature bone tissue formation and greater bone–material affinity. These results demonstrate that electrical surface polarization effectively modulates osteoclast–material interactions and enhances bone regeneration, highlighting its potential as a simple and translatable functionalization strategy for apatite-based bone graft materials.

## 1. Introduction

Bone graft biomaterials play a critical role in bone regeneration. Autografts are widely regarded as the most effective material for bone regeneration due to their excellent osteogenic potential, osteoinductivity, and biocompatibility, as they provide living cells and native growth factors that directly support new bone formation [1]. However, their clinical application is limited by several disadvantages, including donor site morbidity, limited availability of graft material, prolonged surgical time, and increased patient discomfort [1,2]. These limitations have driven the development and extensive investigation of alternative bone graft substitutes such as allografts (human resource), xenografts (animals’ resource) and alloplastic (synthesized) biomaterials [1,3].

Xenografts, derived from natural bone sources, offer a highly porous structure and mineral composition similar to human bone, promoting osteoconduction and structural support for tissue regeneration [4]. For example, Bio-oss® is a widely used xenogeneic bone substitute derived from deproteinized bovine bone mineral and has been extensively applied in clinical bone regeneration procedures due to its excellent biocompatibility and osteoconductivity [5,6]. In the point of view of materials science, Bio-oss® is calcium deficient carbonate apatite with 75-80% porous structures [5–7]. Its natural porous structure closely resembles human cancellous bone, providing an ideal scaffold for cell attachment, vascular infiltration, and new bone formation [8,9]. Numerous clinical and preclinical studies have demonstrated its ability to support long-term bone stability and integration with host tissue [10].

In contrast, alloplastic biomaterials are synthetically engineered, allowing precise control over composition, crystallinity, and surface properties, which enhances reproducibility and enables tailored biological responses [11]. The combined study of xenograft and alloplastic materials provides valuable insights into both clinically relevant and customizable regenerative strategies, supporting the advancement of effective and accessible bone substitute technologies. Despite its favorable biological performance, Bio-Oss® exhibits limited bioactivity and slow resorption rates, which may restrict complete replacement by newly formed bone [12]. Therefore, surface modification and functionalization strategies have been explored to enhance its biological interactions and regenerative capacity and improve bone healing ability.

Surface modification and functionalization strategies have been extensively explored to enhance the osteoconductivity of biomaterials by improving their interactions with cells and surrounding tissues [13,14]. Among these approaches, electrical polarization has emerged as a promising technique to modify surface charge characteristics without altering the bulk properties of the material [15–20]. By inducing permanent or semi-permanent surface charges through the application of an external electric field at elevated temperatures, polarized biomaterials can influence the interactions with biomolecules, cells and surrounding tissues [15–20]. Previous studies have demonstrated that electrically polarized surfaces can promote osteoblast attachment [21–24], proliferation [22], and differentiation, as well as enhance apatite formation [23,25], thereby accelerating bone regeneration [25]. This non-chemical and stable modification method offers a simple effective strategy to improve the osteoconductivity of bone substitute materials and supports the development of advanced functional biomaterials for regenerative applications.

In this study, carbonate apatite (CA) was selected as the model biomaterial for in vitro cell culture evaluation due to its close compositional similarity to the mineral phase of natural bone and its suitability for controlled surface modification, allowing detailed investigation of cellular responses to surface characteristics. This simplified and biomimetic system enables clear assessment of cell adhesion, proliferation, and osteogenic behavior under well-defined conditions. In contrast, a xenograft material was employed for the in vivo animal experiments because it is a widely used and clinically established bone substitute with proven biocompatibility and osteoconductive properties. Utilizing a clinically relevant material for in vivo evaluation enhances the translational significance of the findings and allows assessment of the surface modification strategy in a complex physiological environment. Together, this combined approach integrates mechanistic in vitro insights with practical in vivo validation for bone regeneration applications.

## 2. Materials and Methods

### 2.1. Preparation of biomaterials

CA was synthesized using the procedure described by Doi, et al [26], with slight modifications. CA powder was synthesized using the wet method from analytical grade calcium nitrate tetrahydrate (Kanto-Kagaku, Tokyo, Japan), disodium hydrogen phosphate (Kanto-Kagaku, Tokyo, Japan), and sodium carbonate (Kanto-Kagaku, Tokyo, Japan) reagents with a CO_3_/PO_4_ molar ratio of 5. The CA compacts were pressed in a mold at 200 MPa, and sintered for 2 h in a carbon dioxide atmosphere to minimize carbonate loss from the CA surface at 780 °C. After polishing with a 5 μm grain size diamond disk, the CAs (ϕ7 mm) were washed in ethanol with ultrasonication for 15 min. The specimens were prepared with a thickness of 0.8 mm. The roughness of the specimens was measured using a color laser microscope (Model: OLS4100, Olympus, Japan). Specimens with similar surface roughness (Ra = 0.8 μm) were used in the experiments.

Xenograft materials were prepared from bovine bones. The cortical bovine bones obtained from diaphysis of femur were sliced transversely with a diamond saw to a thickness of approximately 1.0 mm. In the manufacture process, all organic component of Bio-Oss® is removed by heating process (≥16 hours at ≥300°C), followed by alkali treatment (NaOH, pH ≥ 12), and dry-heat sterilization (3 hours at 160°C) to remove proteins from the bovine bones [7]. The bone slices were treated with calcination at 300° C for 24 h, 2M NaOH for 12 h, and heating at 160° C for 3 h. The calcined bone slices were cut in dimensions of 3 × 4 × 0.8 mm. and washed in ethanol with ultrasonication for 15 min.

The CA and xenograft specimens were subjected to material characterization. The XRD measurements were performed for phase analysis at room temperature (RT) with CuKα radiation at 40 kV and 40 mA on a diffraction spectrometer (Model PW1700, Philips, Netherlands). The specimens were subjected to attenuated total reflectance (ATR)-FTIR spectroscopy using a Perkin-Elmer spectrum BS spotlight spectrophotometer with a diamond ATR attachment. The carbonate contents were estimated from the ATR-FTIR spectra using the method described by the ratio of the integrated areas of carbonate-to-phosphate [27,28]. The type-B CA with 8% wt. of carbonate in hydroxyapatite crystal lattice was used in this study.

### 2.2 Electrical polarization treatment

The CA and xenograft specimens were electrically polarized as described previously [29] with a pair of platinum electrodes at 150 −350 °C in direct-current (dc) electric fields of 5 kV·cm^−1^ for 30 min in air (**Table 1**).

**Table 1.** Electrical properties of synthesized CA and xenograft materials.

| Specimen | Polarization | Q (C/cm<sup>2</sup>) |
| --- | --- | --- |
| Synthesized CA | Without polarization | $3 \times 10^{-5}$ |
| | Low (150 °C, 5kV/cm, 30min) | $1 \times 10^{-3}$ |
| | Moderate (250 °C, 5kV/cm, 30min) | $3 \times 10^{-3}$ |
| | High (350 °C, 5kV/cm, 30min) | $11 \times 10^{-3}$ |
| Xenografts | Without polarization | $1.2 \times 10^{-6}$ |
| | Low (150 °C, 5kV/cm, 30min) | $0.51 \times 10^{-3}$ |
| | High (350 °C, 5kV/cm, 30min) | $5 \times 10^{-3}$ |

The thermally stimulated depolarization current (TSDC) was used to measure the electrical energy stored in the bone tissue. TSDC measurements were carried out by previously described method [29] whereby, measurements were carried out in air, at temperatures ranging from RT to 600 (for CA) or 700 (for xenograft materials)°C with a thermal ramp rate of 5.0°C·min^−1^. The depolarization current was measured using a Hewlett–Packard 4140B pA meter (CA, USA). Electrical energy *Q*, activation energy *H*, and relaxation time τ were calculated from the TSDC profiles. *Q* was calculated using equation (1):

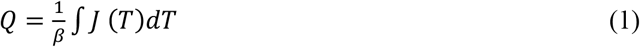

where, *J* (*T*) is the measured dissipation current density at temperature *T*, and β is the heating rate. *Q* and *E*_a_ for depolarization were evaluated from the TSDC profiles using equations (2) and (3), and τ at 37°C, which governs the relaxation process, was calculated using the Arrhenius law (4):

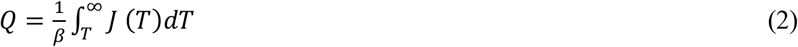

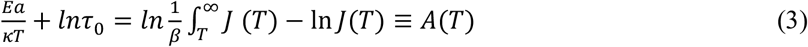

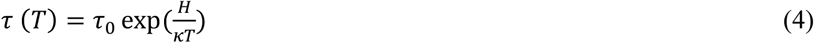

where, J (*T*) is the measured dissipation current density at temperature *T*, β is the heating rate, *τ*_0_ is a pre-exponential factor, and κ is the Boltzmann constant.

### 2.3. Osteoclast culture

Peripheral mononuclear blood cells (PB) were purified from patients as previously described [29]. The donation protocol was approved by the Human Subjects Committee of the University of Turku, Finland (42 /1801/2019). The guidelines in the Declaration of Helsinki were followed, and all patients signed the informed consent paperwork. Briefly, heparin anti-coagulated human blood was diluted 1:1 (v/v) in PBS. Buffy coats layered over Ficoll-Paque solution (GE Healthcare, Little Chalfont, UK) were collected, washed with PBS twice and resuspended in α-MEM (Sigma-Aldrich, MO, USA) supplemented with 10% FBS (Gibco, MA, USA), L-glutamine (Invitrogen, MA, USA), 100 IU/mL penicillin (Invitrogen, MA, USA), and 100 µg/mL streptomycin (Invitrogen, MA, USA). The enriched mononuclear cells were seeded onto the specimens with or without electrical energy at a density of 1×10^6^ cellscm^−2^ and cultured with 20 ng/mL RANKL (PeproTech EC, London, UK) and 10 ng/mL M-CSF (R&D Systems, MN, USA) under humidified atmosphere of 5% CO_2_ at 37°C for 14 days. Half of the medium was changed every 3-4 days.

After the culture, the specimens were delicately scrubbed with brush and washed with distilled water. The dried bone slices were examined under a color laser microscope (Model OLS4100, Olympus, Japan) equipped with an image analysis system. The depth and volume of resorption pits per unit area were measured to quantify the resorption efficacy of the osteoclasts. The measurements were performed for 5 randomly selected areas on 6 slices to obtain an average for each experiment.

### 2.4. Animal study

To study the effects of electrical energy on bone formation, the xenograft materials were implanted into rat femurs [30]. The specimens with or without electrical energy were sterilized using 70% ethanol. The following implantation experiments were carefully completed by veterinarians in accordance with the Guidelines for Animal Experimentation (Tokyo Medical and Dental University, Japan, A2017-099A), as well as the Guide for the Care and Use of Laboratory Animals (National Institutes of Health Pub. No. 85−23, Rev. 1985).

The specimens were surgically implanted into the bilateral femoral diaphysis of Wistar rats (13-week-old males, 400-430g) under isoflurane-based anesthesia. An incision was made in the skin, followed by intermuscular dissection to expose the femoral periosteum, and rectangular defects were created from the cortex to the marrow using a dental engine bur (0.8 mm diameter). Thereafter the bone defects were washed with saline, and the specimens were implanted and fixed in place into the holes by suturing the muscle and skin tissues. After implantation surgery, 1 mg/kg (body weight) butorphanol and 30 mg/kg (body weight) trimethoprim sulfadiazine were injected. After 7 days of the implantation surgery, the rats were sacrificed under anesthesia and perfusion fixed with 4% paraformaldehyde. The femurs were extracted and immersed in 4% paraformaldehyde for 3 days at 4°C. After fixation, each sample was decalcified with 10% neutralized ethylenediamine tetraacetic acid (EDTA) for 5 weeks, dehydrated in an ascending gradient of ethanol concentrations, cleared in xylene, embedded in paraffin, and cut into 5-μm sections. Each section was deparaffinized with xylene, rehydrated with ethanol, and stained with hematoxylin and eosin using standard methods. The affinity index was measured using Image J^®^ software to evaluate the direct bonding between biomaterials and bone tissues. Each experimental group consisted of 5-6 rat femurs for each specimen in a single experiment. The same experiment was repeated three times.

### 2.5. Statistical analysis

Accurate quantifications of the different samples were achieved by performing four independent experiments. Statistical analysis across the experimental groups was performed using the analysis of variance (ANOVA) with Tukey’s post-hoc test for multiple comparisons, using the SPSS software package (version 22, Chicago, IL). The statistically significant level was set at *p* < 0.05 for all the tests. All data has been expressed as mean ± standard deviation (SD).

## 3. Results

The crystalline structure of synthesized CA, natural bone, and bone samples after the alkaline treatment and calcification was analyzed by XRD (**Figure 1**). All samples were highly consistent with published data for hydroxyapatite (ICDD No. 9-432). The CA sample exhibited characteristic apatite diffraction peaks, including prominent reflections corresponding to the 25° (002) and 32° (211) planes, indicating a well-defined apatite phase. In contrast, the bone sample without the alkaline treatment and calcification showed broader peaks, reflecting its lower crystallinity. After the alkaline treatment and calcification, the bone sample demonstrated enhanced apatite-related diffraction features, suggesting improved mineral organization and similarity to carbonate apatite. These results confirm the successful preparation of apatite-based biomaterials with bone-like mineral characteristics.

**Figure 1.**
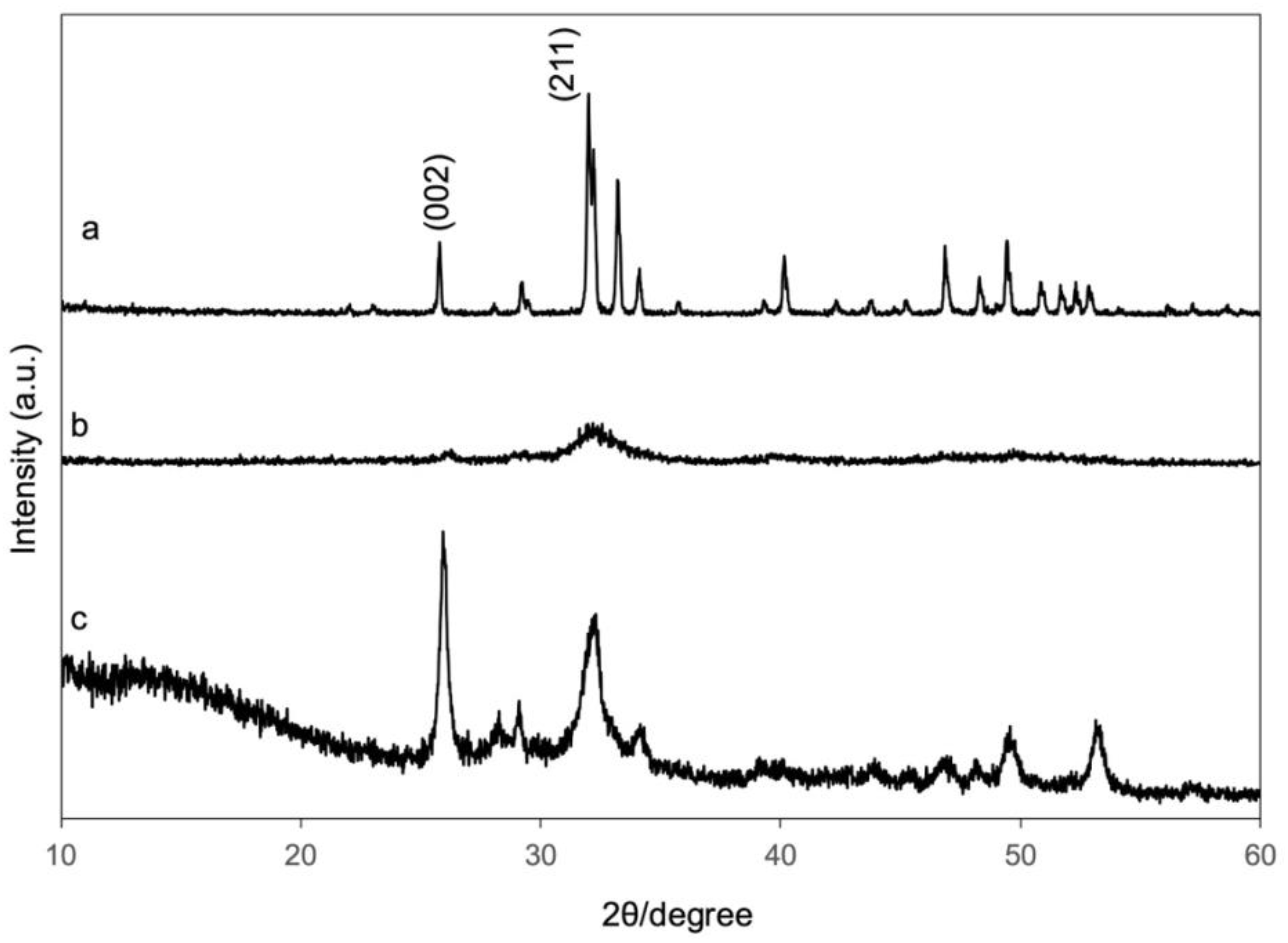
XRD patterns of (a) the synthesized CA, (b) natural bone, and (c) bone samples after alkaline processing and calcification. All samples correspond to the hydroxyapatite phase. The CA sample shows clear apatite peaks at approximately 25° and 32°, while the natural bone exhibits broader peaks indicating lower crystallinity. The treated bone sample demonstrates enhanced apatite features, suggesting improved mineral structure similar to carbonate apatite.

The ATR-FTIR further confirmed the chemical composition of the synthesized CA and bone samples after the alkaline treatment and calcification (**Figure 2**). The ATR-FTIR spectra of all specimens exhibited peaks for phosphate ion absorption at 567, 575, 601,1045, and 1189 cm^−1^, respectively, and carbonate absorption at 871, 1415 and 1450 cm^−1^. Both CA and treated bone samples displayed characteristic phosphate (PO_4_^3−^) absorption bands as well as carbonate (CO_3_^2−^) groups, which are indicative of carbonate-substituted apatite similar to native bone mineral. The presence of these functional groups suggests successful formation of biomimetic apatite structures following treatment, whereas untreated bone exhibited comparatively weaker carbonate-related peaks.

**Figure 2.**
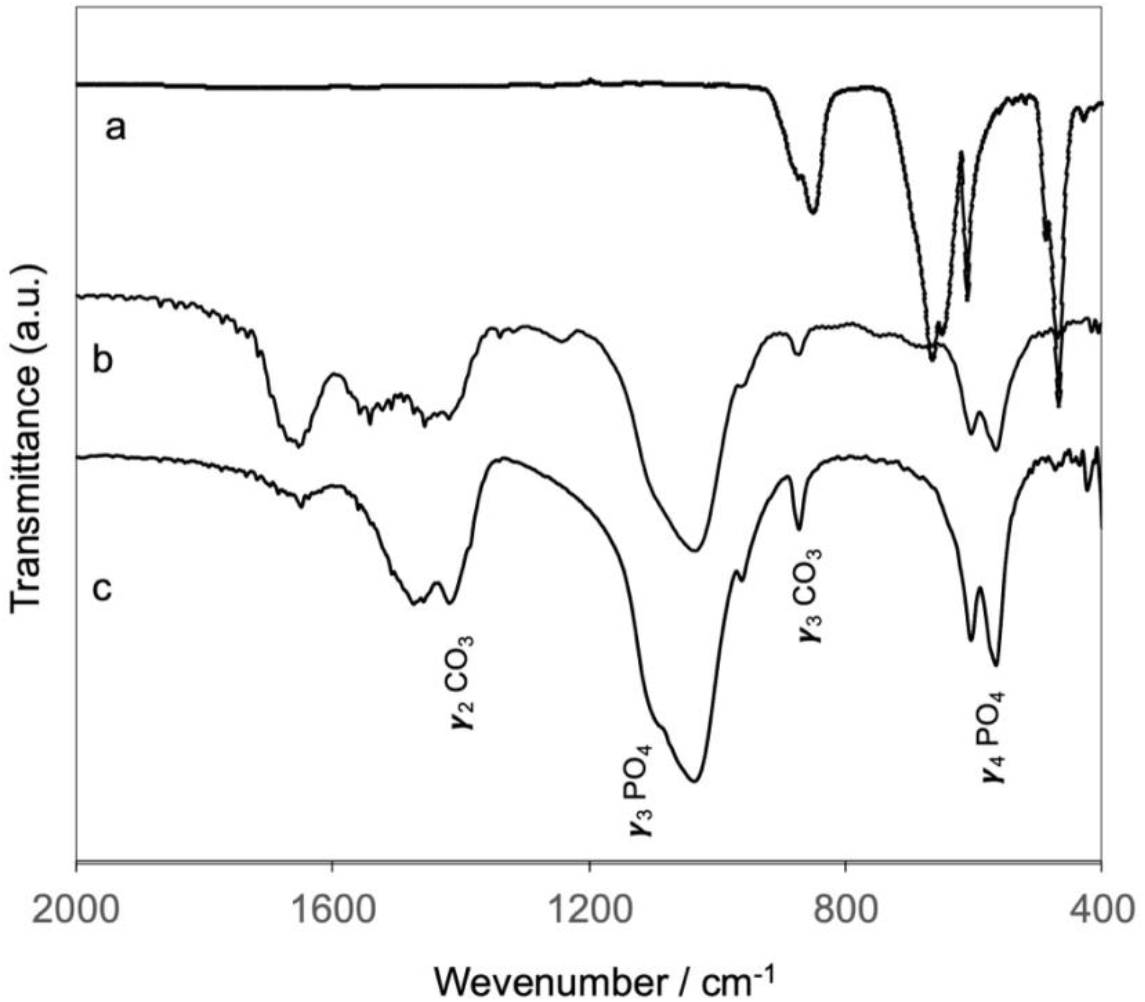
ATR-FTIR spectra of (a) synthesized CA, (b) natural bone, and (c) bone samples after alkaline treatment and calcification. All samples show characteristic phosphate (PO_4_^3−^) bands and carbonate (CO_3_^2−^) bands, indicating carbonate-substituted apatite. The natural bone sample exhibits additional amide bands associated with proteins, which are absent after treatment. The carbonate content of the samples was approximately 8–9 wt%, confirming the formation of B-type carbonate apatite.

The spectra of the bone sample without the alkaline treatment and calcification additionally indicated the adsorption peaks for amide groups adsorption at 1625 (amide I), 1528 (amide II), and 1260 (amide III) cm^−1^, which were characteristics of proteins. It was confirmed that proteins were removed after the alkaline treatment and calcification. The content of carbonate ions in the specimens was calculated from the results of ATR-FTIR was 8.9 wt % for the synthesized CA and 8.0 % for bone after alkaline treatment and calcification. Combined with the results of XRD and ATR-FTIR, the results indicated that the specimens were B-type CA containing approximately 8-9 wt % of carbonate substituted for phosphate sites of the HA lattice.

Typical TSDC curves for the synthesized CA and bone after the alkaline treatment and calcification were shown in **Figure 3a and 3b**, respectively. Polarized samples exhibited increased current density responses compared to non-polarized controls, confirming successful induction of surface charges. The magnitude of polarization increased with higher treatment conditions, indicating that electrical polarization effectively modified the surface electrical properties of the apatite-based materials (**Table 1**).

**Figure 3.**
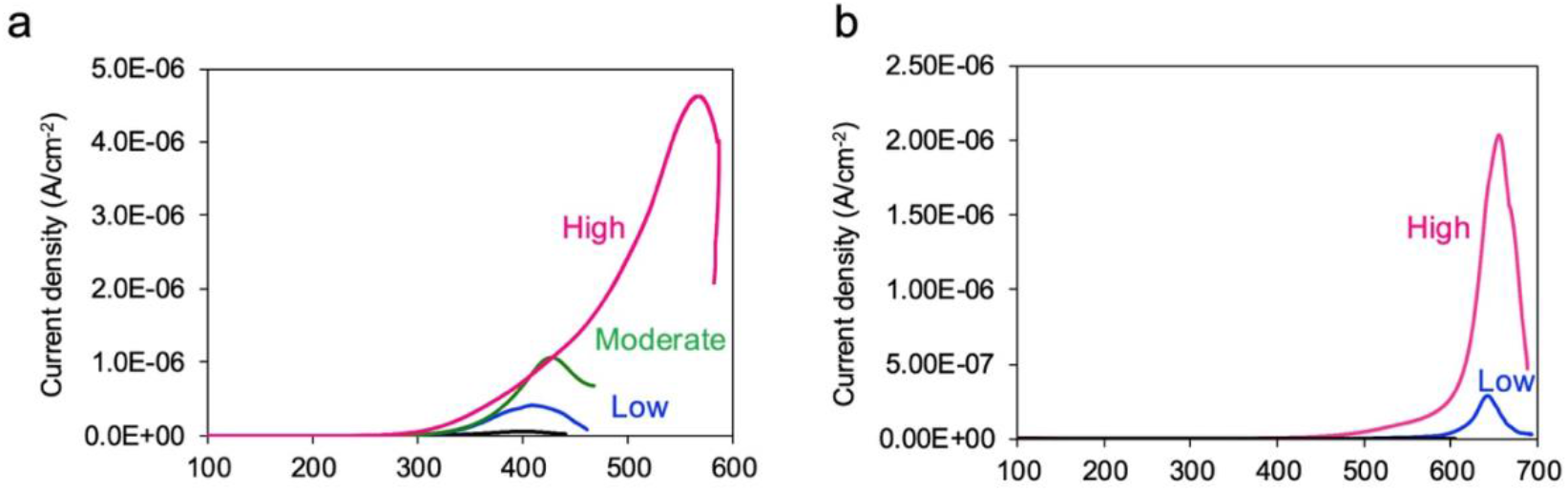
TSDC curves of (a) synthesized CA and (b) bone samples after alkaline treatment and calcification. The CA and xenograft specimens were electrically polarized with a pair of platinum electrodes at 150 (low), 250 (moderate), and 350 (high) °C in dc electric fields of 5 kV·cm^−1^ for 30 min in air. The polarized specimens exhibit higher current density responses compared to non-polarized controls (black line). Increased polarization intensity is observed with higher treatment conditions, indicating effective modification of the surface electrical properties of the apatite-based materials.

In vitro osteoclastic differentiation was assessed using TRAP staining (**Figure 4a**). Representative micrographs revealed the osteoclast precursors differentiated into TRAP-positive multi-nuclear osteoclasts. The osteoclasts cultured on the CA samples are bigger than the ones on bone slices as a control. Quantitative analysis confirmed a statistically significant increase in osteoclast formation on moderately and highly polarized samples, with the greatest enhancement observed on positively polarized surfaces (**Figure 4b**). This result suggests that surface charge induction significantly influences cell–material interactions, likely through modulation of protein adsorption and ionic microenvironments at the material interface. Furthermore, the dependence of cellular response on polarization intensity indicates that surface electrical properties play a critical role in regulating osteoclastic activity, which is essential for balanced bone remodeling processes.

**Figure 4.**
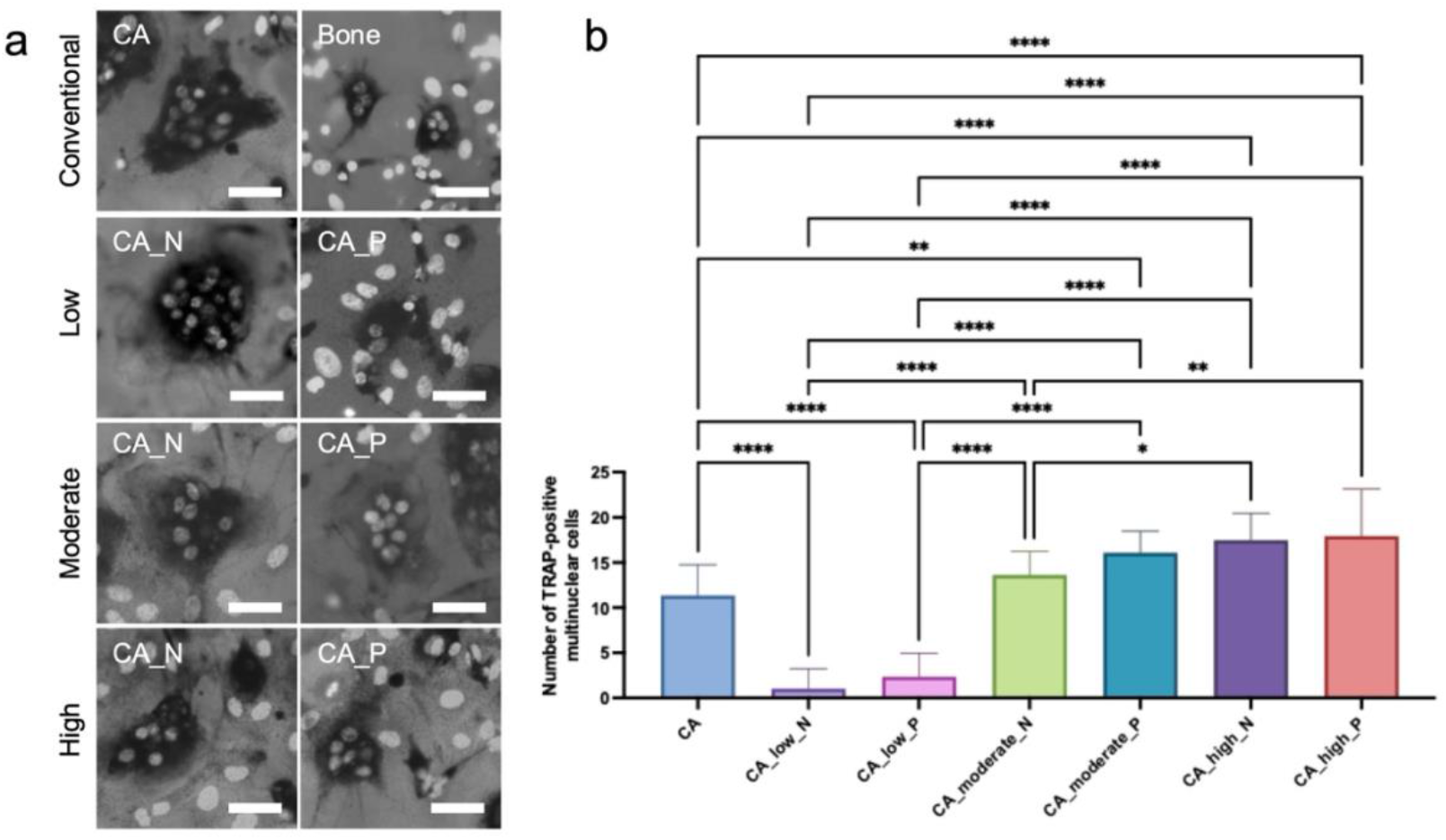
In vitro evaluation of osteoclast differentiation on CA surfaces with different polarization conditions. (a) Representative TRAP-stained micrographs showing multinucleated TRAP-positive osteoclast-like cells on conventional CA and electrically polarized CA surfaces under low, moderate, and high treatment conditions, including negatively (N) and positively (P) polarized samples. Scale bar=100µm (b) Quantitative analysis of TRAP-positive multinucleated cell numbers. Polarized CA surfaces exhibit significantly increased osteoclast formation compared to non-polarized controls, with the highest activity observed on positively polarized and highly polarized samples, indicating that surface charge induction enhances osteoclastic differentiation. Data are shown as the mean ± SD (n=15). \**p* < 0.05, \*\**p* < 0.005, *** *p* < 0.002, \*\*\*\**p* < 0.0001 compared with the conventional CA.

Three-dimensional confocal laser scanning microscopy and quantitative analysis of osteoclast-mediated resorption pits are shown in **Figure 5**. The morphology of the resorption pits made on bone slice as a control show typical shape, indicating that the osteoclast culture system works well. Electrically polarized CA samples exhibited markedly increased resorption activity compared to non-polarized CA (**Figure 5a**). The reconstructed 3D images revealed deeper and more extensive resorption pits on polarized surfaces, particularly in the moderately and highly polarized groups. Quantitative evaluation demonstrated significant increases in both pit depth (**Figure 5b**) and resorbed volume (**Figure 5c**), with the greatest resorption observed on positively polarized CA surfaces. These results indicate that electrical polarization enhances osteoclast functional activity, likely by modifying surface charge characteristics that promote cell adhesion and localized ionic interactions at the material interface. The polarization-dependent increase in resorption further suggests that surface electrical properties play an important role in regulating osteoclast–material interactions.

**Figure 5.**
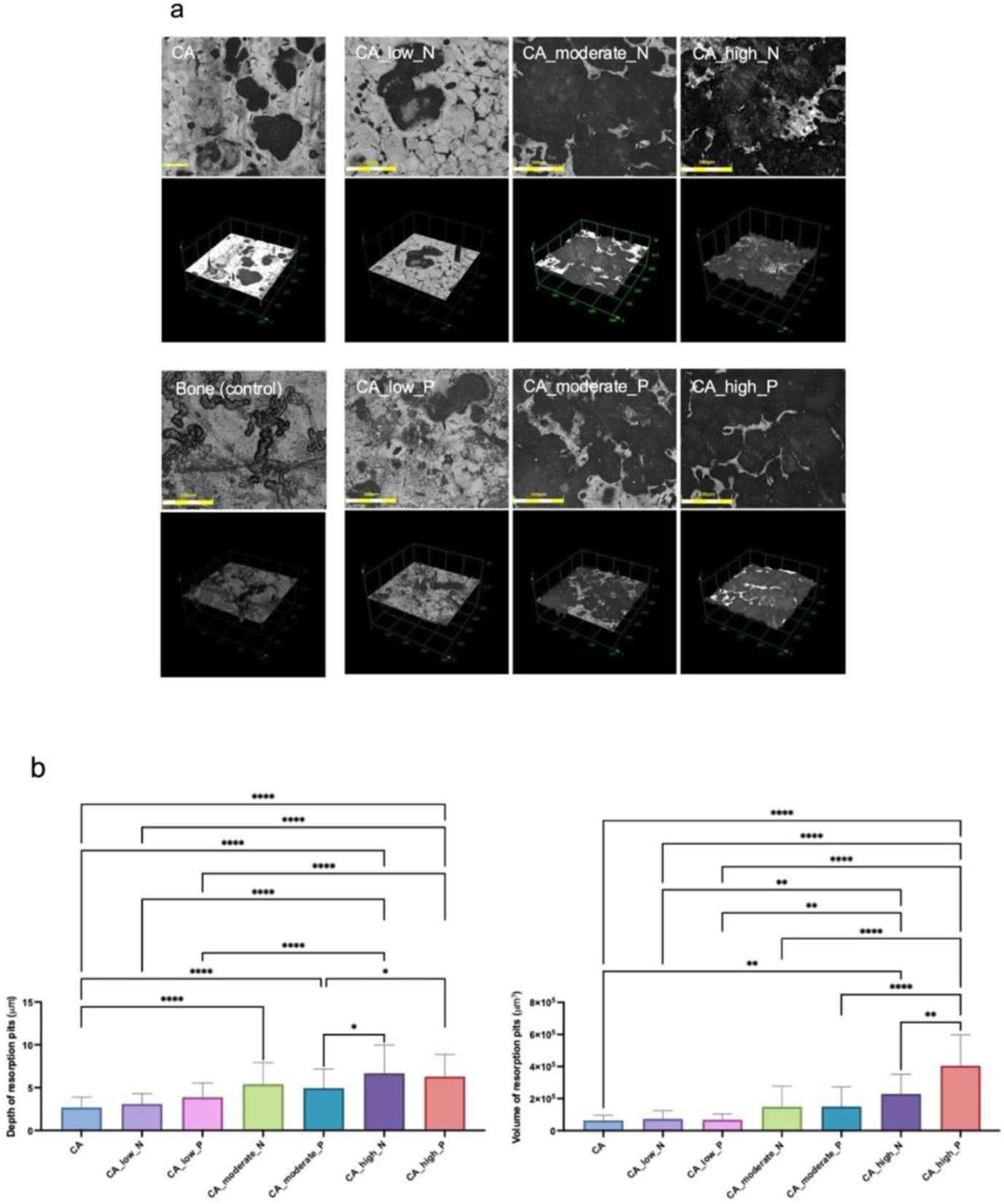
Osteoclast-mediated resorption pit formation on CA surfaces with different polarization conditions. (a) Representative three-dimensional confocal laser scanning microscopy images showing resorption pits on non-polarized CA, natural bone control, and electrically polarized CA surfaces under low, moderate, and high treatment conditions, including negatively (N) and positively (P) polarized samples. Scale bar=200µm (b) Quantitative analysis of resorption pit depth and resorbed volume. Electrically polarized CA surfaces exhibit significantly enhanced resorption activity compared to non-polarized controls, with the greatest pit formation observed on moderately and highly polarized, particularly positively polarized, samples, indicating that surface charge induction promotes osteoclast functional activity. Data are shown as the mean ± SD (n=30-60 for depth measurements and n=12-20 for volume measurements). \**p* < 0.02, \*\**p* < 0.005, *** *p* < 0.002, \*\*\*\**p* < 0.0001 compared with the conventional CA.

In vivo bone regenerative performance was further evaluated through histological examination following implantation of polarized xenograft materials (**Figure 6a**). Histological sections stained with hematoxylin and eosin revealed pronounced new bone formation surrounding polarized implants, with more extensive trabecular structures and reduced fibrous tissue compared to non-polarized controls (**Figure 6b**). Notably, the highly polarized groups exhibited more mature bone tissue with improved osseointegration at the implant–bone interface, indicating accelerated healing and remodeling. Quantitative assessment of affinity index percentage demonstrated significantly higher bone regeneration in samples subjected to higher polarization conditions, particularly on positively charged surfaces (**Figure 6c**). Collectively, these in vivo results confirm that electrical surface polarization effectively enhances osteogenesis and implant integration, highlighting its potential as a functionalization strategy to improve the biological performance of bone graft biomaterials.

**Figure 6.**
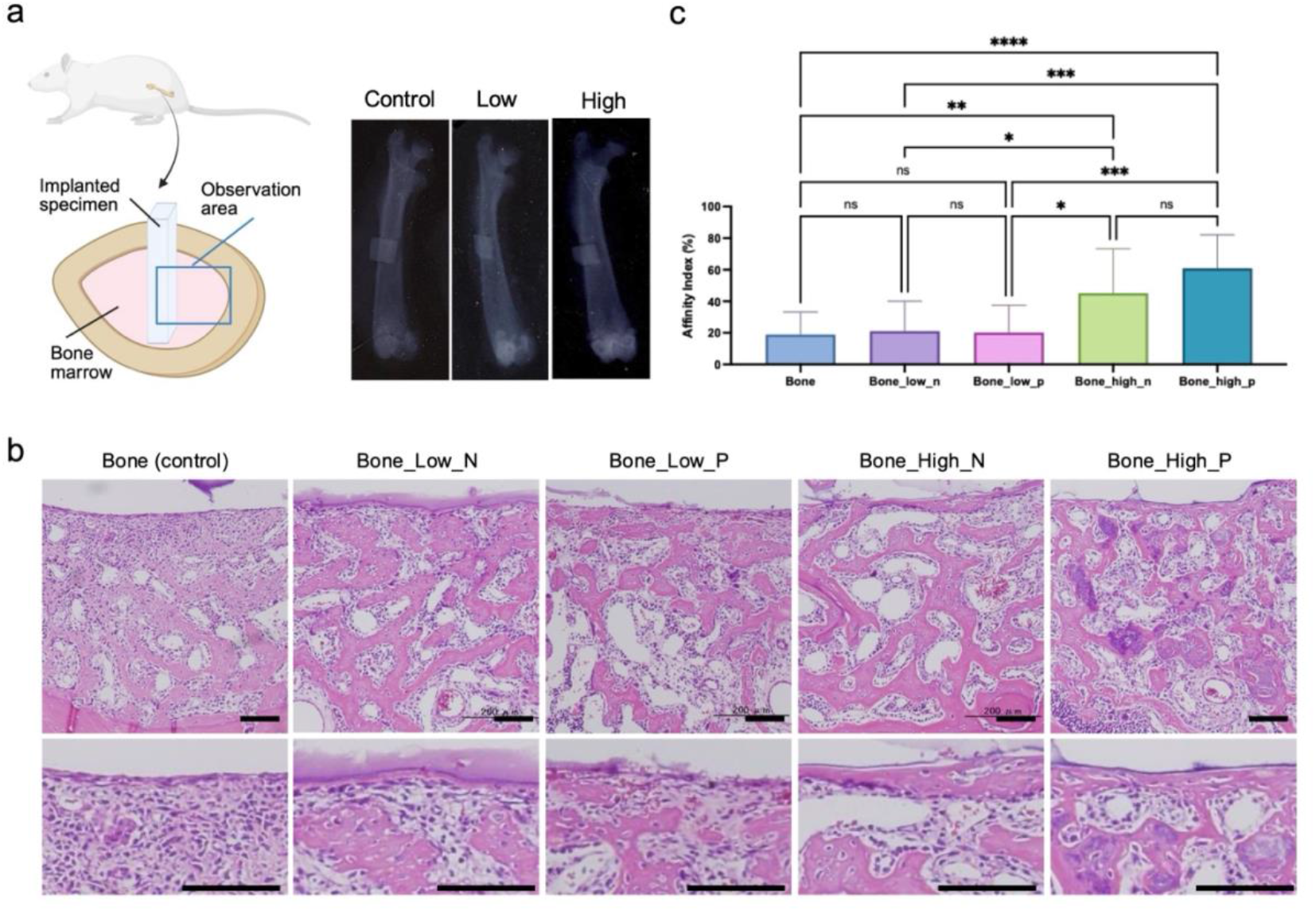
In vivo evaluation of bone regeneration following implantation of non-polarized and electrically polarized xenograft materials. (a) Radiographic images demonstrate the site of bone defects and implantation of materials. The illustration demonstrates the observation area for histology. (b) Representative histological sections stained with hematoxylin and eosin showing new bone formation around implants under different polarization conditions, including low, moderate, and high treatments with negatively (N) and positively (P) polarized surfaces. Enhanced trabecular bone formation and improved osseointegration are observed in polarized groups, particularly in highly polarized samples. Scale bar=100µm. (c) Quantitative analysis of new bone area percentage contacting to the implanted materials, showing significantly higher bone regeneration in polarized samples, with the greatest increase observed in highly and positively polarized groups. Data are shown as the mean ± SD (n=12-36). \**p* < 0.02, \*\**p* < 0.005, *** *p* < 0.002, \*\*\*\**p* < 0.0001 compared with the conventional non-polarized material.

## 4. Discussion

The present study investigated the effect of electrical polarization as a surface functionalization strategy for apatite-based biomaterials, using synthesized CA for in vitro cell evaluation and bone samples after alkaline treatment and calcification as a clinically relevant xenograft for in vivo validation. The results demonstrate that electrical polarization effectively modifies the surface electrical properties of both synthetic and natural apatite materials and significantly enhances osteoclastic activity and bone regeneration.

Material characterization by XRD and ATR-FTIR confirmed that both synthesized CA and bone samples after alkaline treatment and calcification possessed crystalline structures and chemical compositions consistent with carbonate-substituted hydroxyapatite (**Figure 1 and 2**) bone [31,32]. The presence of characteristic phosphate and carbonate bands, along with the calculated carbonate content of approximately 8–9 wt%, indicates the B-type carbonate apatite, which closely resembles the mineral phase of natural bone [31,32]. The removal of organic components from the bone matrix after treatment was further verified by the disappearance of amide-related peaks in the ATR-FTIR spectra. These results suggest that the prepared biomaterials successfully mimicked native bone mineral, providing a suitable material for investigating the effects of surface polarization on biological responses.

The TSDC measurements demonstrated successful induction of surface charges on both CA and treated bone samples, with polarization intensity increasing under higher treatment conditions (**Figure 3 and Table 1**). This confirms that electrical polarization is an effective and controllable approach for modifying the surface electrical characteristics of apatite-based biomaterials without altering their bulk structure.

The higher electrical polarization energy observed in the synthesized CA compared to xenograft material can be attributed primarily to differences in their microstructural characteristics. Synthesized CA is a relatively dense ceramic material with low porosity, which allows for more efficient dipole alignment and charge storage within the crystal lattice during the polarization process [29,33]. In contrast, Bio-Oss® is known as a calcium-deficient carbonate apatite with a highly porous structure, typically exhibiting 75–80% interconnected porosity [5–7]. While this porous architecture is advantageous for cell infiltration and vascularization, it reduces the effective volume of solid material available for charge accumulation and dipole orientation [23,34,35]. The presence of extensive pore spaces disrupts continuous crystal domains and limits the overall dielectric capacity of the material, resulting in lower polarization energy and reduced surface charge density. From a materials science perspective, denser structures possess higher dielectric constants and greater capacity to retain induced charges, thereby generating stronger electrical fields at the surface.

In vitro cell evaluation revealed that electrically polarized CA surfaces significantly enhanced osteoclastic differentiation and resorption (**Figure 4 and 5**). These results suggest that surface electrical charges promote osteoclast differentiation and resorption, likely through electrostatic interactions that influence protein adsorption patterns and local ionic environments [36,37]. Enhanced osteoclastic resorption activity is a critical component of bone remodeling, as it facilitates the resorption of biomaterials and creates space for subsequent new bone formation [38,39]. The enhanced osteoclastic differentiation and resorptive activity observed on electrically polarized CA surfaces can be attributed to the modification of surface charge, which strongly influences biomolecular interactions at the material interface.

Polarized surfaces are known to alter the adsorption behavior, orientation, and conformation of serum proteins such as bone serum albumin, fibrinogen, [40], fibronectin, and osteopontin [41], which play essential roles in the adhesion of macrophage as a osteoclast precursor and subsequent cell fusion processes [41]. Improved protein adsorption on charged surfaces provides favorable binding sites for integrin-mediated signaling pathways that regulate osteoclast differentiation and resorption [39]. In addition, surface charges can modify the local ionic microenvironment by attracting calcium and phosphate ions, which are critical regulators of osteoclast function and resorption activity [42–44]. The accumulation of these ions may enhance osteoclast maturation and promote the formation of the acidic microenvironment required for mineral dissolution during resorption activity [42–44].

The polarization-dependent increase in osteoclastic activity further indicates that stronger surface charge densities intensify these biological effects. Higher polarization levels generate more pronounced electrostatic fields, leading to increased protein adsorption and greater ionic accumulation at the interface, thereby amplifying cellular responses [24,33]. This explains the observed enhancement in both osteoclast differentiation and resorption pit formation under moderate and high polarization conditions.

The positively charged surfaces exhibited greater osteoclastic activity compared to negatively charged surfaces. This phenomenon may be explained by the preferential adsorption of negatively charged biomolecules, including phosphate-containing proteins and glycoproteins (e.g. bovine serum albumin and osteopontin), onto positively charged surfaces through electrostatic. Such proteins are known to facilitate osteoclast adhesion, cytoskeletal organization, and activation attraction [40,41]. Furthermore, positively charged surfaces can more effectively attract negatively charged phosphate ions while simultaneously concentrating calcium ions in the surrounding microenvironment, creating favorable conditions for osteoclast-mediated mineral dissolution [44]. Previous studies have also reported enhanced osteoclast and osteoblast responses on positively polarized apatite, supporting the critical role of surface polarity in regulating bone cell behavior [24,29,33].

The in vivo results corroborated the in vitro findings, demonstrating that electrical polarization significantly improved bone regeneration around xenograft materials (**Figure 6**). Histological analysis revealed more extensive new bone formation in polarized groups, particularly under high polarization conditions. Quantitative assessment of bone regeneration showed significantly higher values for polarized xenograft materials, with positively charged surfaces exhibiting the greatest osteogenic response. These observations indicate that electrical polarization not only enhances cellular responses in vitro but also translates into improved regenerative outcomes in a complex physiological environment.

The enhanced bone formation observed in polarized xenograft materials may be attributed to several mechanisms. Surface charges can modulate the adsorption and conformation of serum proteins such as bovine serum albumin and fibronectin, which are essential for cell attachment and signaling [40,41]. Additionally, polarized surfaces may promote the accumulation of calcium and phosphate ions, facilitating mineral deposition and accelerating bone formation [42–44]. The polarization-dependent responses observed in both osteoclastic activity and in vivo bone regeneration further suggest that the magnitude and polarity of surface charges play crucial roles in regulating biological interactions.

While the present study focused on osteoclastic responses and bone regeneration, further investigations into osteoblastic behavior, angiogenesis, and long-term remodeling processes would provide a more comprehensive understanding of the biological effects of polarized surfaces. Additionally, optimization of polarization parameters, including treatment temperature, electric field strength, and charge stability over time, may further enhance regenerative outcomes. Compared with conventional surface modification techniques involving chemical coatings or biomolecule immobilization [14,45], electrical polarization offers a simple, stable, and non-chemical approach to enhance the bioactivity of bone graft materials. This strategy preserves the intrinsic properties of the biomaterials while providing functional surface characteristics that actively promote tissue regeneration. The successful application of polarization to both synthetic CA and clinically used xenograft materials underscores its versatility and translational potential.

## 5. Conclusion

This study demonstrates that electrical surface polarization effectively enhances the biological performance of apatite-based biomaterials by promoting osteoclastic activity and accelerating bone regeneration. By combining mechanistic in vitro evaluation using carbonate apatite with clinically relevant in vivo assessment using xenograft materials, the findings provide strong evidence for the potential of electrical polarization as a functionalization strategy for advanced bone graft substitutes.

## 6. Acknowledgement

This study was supported by the Sigrid Juselius Foundation, JSPS Grants-in-Aid for Scientific Research (JP23K08670), The Murata Foundation and the Turku Collegium for Science and Medicine. We thank Ms. Naoko Hori, Ms. Takako Takuma (Institute of Science Tokyo), Ms. Katja Sampalahti, Ms. Tatiana Peskova, and Dr. Vuokko Loimaranta (University of Turku) for their technical assistance.

## References

[1] Oryan, S. Alidadi, A. Moshiri, N. Maffulli, Bone regenerative medicine: classic options, novel strategies, and future directions., J. Orthop. Surg. Res. 9 (2014) 18. 10.1186/1749-799X-9-18.

[2] R.R. Betz, Limitations of autograft and allograft: new synthetic solutions., Orthopedics. 25 (2002) s561–70. 10.3928/0147-7447-20020502-04.

[3] R.J. Miron, Optimized bone grafting., Periodontol. 2000. 94 (2024) 143–160. 10.1111/prd.12517.

[4] R. Amid, A. Kheiri, L. Kheiri, M. Kadkhodazadeh, M. Ekhlasmandkermani, Structural and chemical features of xenograft bone substitutes: A systematic review of in vitro studies., Biotechnol. Appl. Biochem. 68 (2021) 1432–1452. 10.1002/bab.2065.

[5] M. Figueiredo, J. Henriques, G. Martins, F. Guerra, F. Judas, H. Figueiredo, Physicochemical characterization of biomaterials commonly used in dentistry as bone substitutes--comparison with human bone., J. Biomed. Mater. Res. Part B Appl. Biomater. 92 (2010) 409–419. 10.1002/jbm.b.31529.

[6] T. Accorsi-Mendonça, M.B. Conz, T.C. Barros, L.A. de Sena, G. de A. Soares, J.M. Granjeiro, Physicochemical characterization of two deproteinized bovine xenografts., Braz. Oral Res. 22 (2008) 5–10. 10.1590/s1806-83242008000100002.

[7] S. Kasuya, N. Kato-Kogoe, M. Omori, K. Yamamoto, S. Taguchi, H. Fujita, N. Imagawa, A. Sunano, K. Inoue, Y. Ito, A. Hirata, T. Ueno, P.K. Moy, New Bone Formation Process Using Bio-Oss and Collagen Membrane for Rat Calvarial Bone Defect: Histological Observation., Implant Dent. 27 (2018) 158–164. 10.1097/ID.0000000000000738.

[8] V. Sollazzo, A. Palmieri, L. Scapoli, M. Martinelli, A. Girardi, F. Alviano, A. Pellati, V. Perrotti, F. Carinci, Bio-Oss®acts on Stem cells derived from Peripheral Blood., Oman Med. J. 25 (2010) 26–31. 10.5001/omj.2010.7.

[9] E.S. Tadjoedin, G.L. de Lange, A.L.J.J. Bronckers, D.M. Lyaruu, E.H. Burger, Deproteinized cancellous bovine bone (Bio-Oss) as bone substitute for sinus floor elevation. A retrospective, histomorphometrical study of five cases., J. Clin. Periodontol. 30 (2003) 261–270. 10.1034/j.1600-051x.2003.01099.x.

[10] Rapone, A.D. Inchingolo, S. Trasarti, E. Ferrara, E. Qorri, A. Mancini, N. Montemurro, A. Scarano, A.M. Inchingolo, G. Dipalma, F. Inchingolo, Long-Term Outcomes of Implants Placed in Maxillary Sinus Floor Augmentation with Porous Fluorohydroxyapatite (Algipore® FRIOS®) in Comparison with Anorganic Bovine Bone (Bio-Oss®) and Platelet Rich Plasma (PRP): A Retrospective Study., J. Clin. Med. 11 (2022). 10.3390/jcm11092491.

[11] S.K. Wong, M.M.F. Yee, K.-Y. Chin, S. Ima-Nirwana, A review of the application of natural and synthetic scaffolds in bone regeneration., J. Funct. Biomater. 14 (2023). 10.3390/jfb14050286.

[12] L.F. Gil, V.V. Nayak, E.B. Benalcázar Jalkh, N. Tovar, K.-J. Chiu, J.C. Salas, C. Marin, M. Bowers, G. Freitas, D.C. Mbe Fokam, P.G. Coelho, L. Witek, Laddec® versus Bio-Oss®: The effect on the healing of critical-sized defect - Calvaria rabbit model., J. Biomed. Mater. Res. Part B Appl. Biomater. 110 (2022) 2744–2750. 10.1002/jbm.b.35125.

[13] H. Amani, H. Arzaghi, M. Bayandori, A.S. Dezfuli, H. Pazoki-Toroudi, A. Shafiee, L. Moradi, Controlling Cell Behavior through the Design of Biomaterial Surfaces: A Focus on Surface Modification Techniques, Adv. Mater. Interfaces. 6 (2019) 1900572. 10.1002/admi.201900572.

[14] X. Hu, T. Wang, F. Li, X. Mao, Surface modifications of biomaterials in different applied fields., RSC Adv. 13 (2023) 20495–20511. 10.1039/d3ra02248j.

[15] X. Ni, Y. Cui, M. Salehi, M.L.S. Nai, K. Zhou, C. Vyas, B. Huang, P. Bartolo, Piezoelectric Biomaterials for Bone Regeneration: Roadmap from Dipole to Osteogenesis., Adv Sci (Weinh). (2025) e14969. 10.1002/advs.202414969.

[16] K.K. Das, R. Pandey, A.K. Dubey, Piezo-electronics: A paradigm for self-powered bioelectronics., Biomaterials. 318 (2025) 123118. 10.1016/j.biomaterials.2025.123118.

[17] X. Chen, S. Zhang, S. Peng, Y. Qian, J. Zhou, Piezoelectric materials for bone implants: Opportunities and challenges, Nano Energy. 138 (2025) 110841. 10.1016/j.nanoen.2025.110841.

[18] B.C. Heng, Y. Bai, X. Li, L.W. Lim, W. Li, Z. Ge, X. Zhang, X. Deng, Electroactive Biomaterials for Facilitating Bone Defect Repair under Pathological Conditions., Adv Sci (Weinh). 10 (2023) e2204502. 10.1002/advs.202204502.

[19] A. Bigham, M.G. Raucci, K. Zheng, A.R. Boccaccini, L. Ambrosio, Oxygen-Deficient Bioceramics: Combination of Diagnosis, Therapy, and Regeneration., Adv. Mater. 35 (2023) e2302858. 10.1002/adma.202302858.

[20] J. Sans, M. Arnau, A. Fontana-Escartín, P. Turon, C. Alemán, Permanently Polarized Materials: An Approach for Designing Materials with Customized Electrical Properties, Chem. Mater. 35 (2023) 3765–3780. 10.1021/acs.chemmater.3c00453.

[21] M. Nakamura, A. Nagai, Y. Tanaka, Y. Sekijima, K. Yamashita, Polarized hydroxyapatite promotes spread and motility of osteoblastic cells., J. Biomed. Mater. Res. A. 92 (2010) 783–790. 10.1002/jbm.a.32404.

[22] D. Kumar, J.P. Gittings, I.G. Turner, C.R. Bowen, L.A. Hidalgo-Bastida, S.H. Cartmell, Polarization of hydroxyapatite: influence on osteoblast cell proliferation., Acta Biomater. 6 (2010) 1549–1554. 10.1016/j.actbio.2009.11.008.

[23] S.H. Cartmell, S. Thurstan, J.P. Gittings, S. Griffiths, C.R. Bowen, I.G. Turner, Polarization of porous hydroxyapatite scaffolds: influence on osteoblast cell proliferation and extracellular matrix production., J. Biomed. Mater. Res. A. 102 (2014) 1047–1052. 10.1002/jbm.a.34790.

[24] M. Nakamura, A. Nagai, T. Hentunen, J. Salonen, Y. Sekijima, T. Okura, K. Hashimoto, Y. Toda, H. Monma, K. Yamashita, Surface electric fields increase osteoblast adhesion through improved wettability on hydroxyapatite electret., ACS Appl. Mater. Interfaces. 1 (2009) 2181–2189. 10.1021/am900341v.

[25] T. Kobayashi, S. Itoh, S. Nakamura, M. Nakamura, K. Shinomiya, K. Yamashita, Enhanced bone bonding of hydroxyapatite-coated titanium implants by electrical polarization., J. Biomed. Mater. Res. A. 82 (2007) 145–151. 10.1002/jbm.a.31080.

[26] Y. Doi, T. Koda, N. Wakamatsu, T. Goto, H. Kamemizu, Y. Moriwaki, M. Adachi, Y. Suwa, Influence of carbonate on sintering of apatites., J. Dent. Res. 72 (1993) 1279–1284. 10.1177/00220345930720090401.

[27] A.R. Cassella, R.C. de Campos, S. Garrigues, M. de la Guardia, A. Rossi, Fourier transform infrared determination of CO2 evolved from carbonate in carbonated apatites., Fresenius. J. Anal. Chem. 367 (2000) 556–561. 10.1007/s002160000368.

[28] E.P. Paschalis, E. DiCarlo, F. Betts, P. Sherman, R. Mendelsohn, A.L. Boskey, FTIR microspectroscopic analysis of human osteonal bone., Calcif. Tissue Int. 59 (1996) 480–487. 10.1007/BF00369214.

[29] L. Bergara-Muguruza, K. Mäkelä, T. Yrjälä, J. Salonen, K. Yamashita, M. Nakamura, Surface Electric Fields Increase Human Osteoclast Resorption through Improved Wettability on Carbonate-Incorporated Apatite., ACS Appl. Mater. Interfaces. 13 (2021) 58270–58278. 10.1021/acsami.1c14358.

[30] M. Nakamura, R. Hiratai, T. Hentunen, J. Salonen, K. Yamashita, Hydroxyapatite with High Carbonate Substitutions Promotes Osteoclast Resorption through Osteocyte-like Cells, ACS Biomater. Sci. Eng. 2 (2016) 259–267. 10.1021/acsbiomaterials.5b00509.

[31] R. Zapanta-LeGeros, Effect of carbonate on the lattice parameters of apatite., Nature. 206 (1965) 403–404. 10.1038/206403a0.

[32] C. Rey, V. Renugopalakrishnan, B. Collins, M.J. Glimcher, Fourier transform infrared spectroscopic study of the carbonate ions in bone mineral during aging., Calcif. Tissue Int. 49 (1991) 251–258. 10.1007/BF02556214.

[33] M. Nakamura, N. Hori, H. Ando, S. Namba, T. Toyama, N. Nishimiya, K. Yamashita, Surface free energy predominates in cell adhesion to hydroxyapatite through wettability., Mater. Sci. Eng. C Mater. Biol. Appl. 62 (2016) 283–292. 10.1016/j.msec.2016.01.037.

[34] T. Iwasaki, Y. Tanaka, M. Nakamura, A. Nagai, K. Hashimoto, Y. Toda, K. Katayama, K. Yamashita, Rate of bonelike apatite formation accelerated on polarized porous hydroxyapatite, J. Am. Ceram. Soc. 91 (2008) 3943–3949. 10.1111/j.1551-2916.2008.02766.x.

[35] S. Nakamura, T. Kobayashi, M. Nakamura, S. Itoh, K. Yamashita, Electrostatic surface charge acceleration of bone ingrowth of porous hydroxyapatite/beta-tricalcium phosphate ceramics., J. Biomed. Mater. Res. A. 92 (2010) 267–275. 10.1002/jbm.a.32354.

[36] M. Nakamura, K. Niwa, S. Nakamura, Y. Sekijima, K. Yamashita, Interaction of a blood coagulation factor on electrically polarized hydroxyapatite surfaces., J. Biomed. Mater. Res. Part B Appl. Biomater. 82 (2007) 29–36. 10.1002/jbm.b.30701.

[37] S. Bodhak, S. Bose, A. Bandyopadhyay, Role of surface charge and wettability on early stage mineralization and bone cell-materials interactions of polarized hydroxyapatite., Acta Biomater. 5 (2009) 2178–2188. 10.1016/j.actbio.2009.02.023.

[38] H. Wang, G. Yang, Y. Xiao, G. Luo, G. Li, Z. Li, Friend or foe? essential roles of osteoclast in maintaining skeletal health., Biomed Res. Int. 2020 (2020) 4791786. 10.1155/2020/4791786.

[39] Q. Xiang, L. Li, W. Ji, D. Gawlitta, X.F. Walboomers, J.J.J.P. van den Beucken, Beyond resorption: osteoclasts as drivers of bone formation., Cell Regen (Lond). 13 (2024) 22. 10.1186/s13619-024-00205-x.

[40] P. Tworek, K. Rakowski, M. Szota, M. Lekka, B. Jachimska, Changes in Secondary Structure and Properties of Bovine Serum Albumin as a Result of Interactions with Gold Surface., ChemPhysChem. 25 (2024) e202300505. 10.1002/cphc.202300505.

[41] J. Maciel, M.I. Oliveira, R.M. Gonçalves, M.A. Barbosa, The effect of adsorbed fibronectin and osteopontin on macrophage adhesion and morphology on hydrophilic and hydrophobic model surfaces., Acta Biomater. 8 (2012) 3669–3677. 10.1016/j.actbio.2012.06.010.

[42] I.S. Harding, N. Rashid, K.A. Hing, Surface charge and the effect of excess calcium ions on the hydroxyapatite surface., Biomaterials. 26 (2005) 6818–6826. 10.1016/j.biomaterials.2005.04.060.

[43] M. Zaidi, O.A. Adebanjo, B.S. Moonga, L. Sun, C.L. Huang, Emerging insights into the role of calcium ions in osteoclast regulation., J. Bone Miner. Res. 14 (1999) 669–674. 10.1359/jbmr.1999.14.5.669.

[44] M. Kanatani, T. Sugimoto, J. Kano, M. Kanzawa, K. Chihara, Effect of high phosphate concentration on osteoclast differentiation as well as bone-resorbing activity., J. Cell. Physiol. 196 (2003) 180–189. 10.1002/jcp.10270.

[45] J. Alipal, N.A.S. Mohd Pu’ad, N.H.M. Nayan, N. Sahari, H.Z. Abdullah, M.I. Idris, T.C. Lee, An updated review on surface functionalisation of titanium and its alloys for implants applications, Materials Today: Proceedings. 42 (2021) 270–282. 10.1016/j.matpr.2021.01.499.

